# Non-apoptotic role of Cleaved Caspase 3 in Rat Gonocyte

**DOI:** 10.1101/2020.02.18.953646

**Authors:** Marina Nunes, Anelise Diniz Arantes, Renato Borges Tesser, Priscila Henriques da Silva, Leticia Rocha da Silva, Samara Urban de Oliva, Ana Flávia Popi, Taiza Stumpp

## Abstract

Germ cells emerge from the epiblast and migrate to the gonads, where they become gonocytes. The gonocytes are the precursors of the spermatogonial stem cells, but little is known about their differentiation. The rigid control of gonocyte proliferation, quiescence and pluripotency marker expression is crucial for spermatogonia development. We have previously suggested that cleaved caspase-3 (Casp3) might play a non-apoptotic role in gonocyte quiescence. Here we describe when rat fetal gonocyte enter mitotic arrest and show that Casp3 inhibition in these cells affects the expression of cell cycle genes. The expression of Ki67, p27^Kip^, Retinoblastoma 1 (pRb1), NANOG and CASP3 was investigated in 15, 17 and 19 days post coitum rat embryo gonads. The results show that Ki67 and pRB1 proteins are downregulated from 15 days post coitum to 19 days post coitum, whereas p27^Kip^, NANOG and CASP3 are upregulated. This suggests that rat germ cells start to enter quiescence around 15dpc and that CASP3 and NANOG seem to play a role in this process. CASP3 labelling formed a ring in gonocyte cytoplasm, which is clearly distinct from apoptotic cell labelling, and coincided with NANOG labelling. CASP3 inhibition lead to an increase of *Pcna* expression and to a decrease of *p27*^*kip*^ and *p21*^*cip*^ expression. These results suggest that cleaved CASP3 has a role in rat male germ cell development which can be related to the control of the cell cycle genes.

## Introduction

Male germ cells reach complete development in the peripubertal phase of life in mammals. Their development begins very early in the embryonic life and involves complex processes such as the inhibition of the somatic program, definition of the germ cell fate and sex differentiation in the gonads [1,2].

In mice, primordial germ cells (PGC) derive from a pool of cells from the posterior epiblast [3] from where they migrate through the hindgut and its mesentery to reach the gonad ridge [4]. In the gonads the germ cells proliferate for a short period of time and are committed to male or female differentiation. Female germ cells seem to follow an intrinsic program and enter meiosis [5]. The male differentiation programme involves at least two critical phenomena: Inhibition of meiosis and a quiescent period. In the early embryonic testes Sertoli cells seem to be responsible for inhibiting meiosis and for inducing male germ cell differentiation [6], although germ cell also seems to have a role in testis differentiation [7]. Male germ cell quiescence also seems to be induced by the Sertoli cells through physical interactions mediated by Kit receptor and stem cell factor (SCF) [8] and via secretion of growth factors such as Activin A that has inhibitory effects on cell proliferation in foetal rat testis organ cultures [9].

The significance of the quiescence period to male germ cell development has not been unveiled yet, although it is known that during this period gonocytes undergo key processes such as redistribution along the seminiferous cords [10] and migration to the basis of the seminiferous epithelium via c-Kit/SCF interaction to differentiate into spermatogonia [6, 11, 12]. In the basis of the seminiferous epithelium gonocytes exit quiescence, what also seems to be regulated by Sertoli cells via NOTCH1 signaling [13] and differentiate into the first spermatogonia [11]. Thus, it is clear that the quiescence is not a period of cell inactivity, although the mechanisms involved in this phenomenon are poorly understood.

The quiescence period of the gonocytes can be considered a particular kind of cell cycle interruption, since it is not related to terminal differentiation or senescence. Instead, it precedes the appearance of the male germ stem cells. In mice, male gonocytes enter the G0 stage between 12.5 and 14.5 days post-coitum (dpc) through Rb1 dephosphorylation and activation of p27^kip^, p15^INK4b^ and p16^INK4a^ and p21^Cip^ [14]. In rats, Beaumont and Mandl (1963) showed that gonocytes do not proliferate at 18.5dpc [15]. However, a description of the time-point when rat gonocytes enter quiescence and of the proteins involved is missing. Previous data obtained by our group suggested that the cleaved Caspase-3 (CASP3) seems to have a non-apoptotic function during rat gonocyte development [10]. Caspases are cysteine-proteases that cleave many different proteins at specific sites [16-19]. Caspase substrates include transcription regulation and cell cycle proteins, such as p27^kip^ [20]. It has been suggested that CASP3 has a role in the control of cell cycle regulation through the processing of RB1 [21] and p21^cip^ [22] indicating that CASP3 has other functions but driving apoptosis. In mice, RB1 and p27^kip^ have important roles in the control of male germ cell cycle and are fundamental for gonocyte quiescence [14, 23].

Here we used cell cycle markers to investigate at what age rat male germ cells enter mitotic arrest and reach full quiescence and suggest that cleaved CASP3 has a role in this process.

## Materials and methods

### Animals and Tissue Preparation

Male and female adult rats (*Rattus norvegicus albinus*) were obtained from the Centre for Development of Experimental Models for Medicine and Biology (CEDEME) of the Paulista Medicine School (EPM) of the Federal University of Sao Paulo (UNIFESP). The matings were performed in the Laboratory of Developmental Biology of the Department of Morphology and Genetics (EPM/UNIFESP, Sao Paulo – Brazil). The animals were kept in plastic cages under a 12–12 hours light/dark cycle at 23-25° C. Food and water were allowed *ad libitum*. Pregnancy was detected by the presence of sperm in the vaginal smears (1dpc).

### Embryo collection

The dams were euthanized by analgesic/anaesthetic (xylazin/ketamin, 10 mg/Kg and 100 mg/Kg, respectively) method. The embryos were collected at 15, 17 and 19dpc and fixed in Bouin’s solution or Carnoy’s solution for immunohistochemistry. Embryo sexing was performed by visual inspection of the gonads. The embryos from each age were obtained from five different mothers to guarantee sample variability and five embryos were used for each analysis. The male gonads were submitted to Ki67, pRB1, PCNA, p27^kip^, NANOG and CASP3 labelling, as described later.

For subsequent analysis, 19dpc embryos were collected from 9 females and had their testes collected for *in vitro* experiment (item 2.4) and RT-qPCR. Ten to twelve embryos were used for this experiment. Finally, 19dpc embryos from other 3 pregnant females were used for Western blot analysis (item 2.6).

### Immunohistochemistry

Six embryos (15 and 17dpc) or testes (19dpc) were fixed in Bouin’s liquid and processed for paraffin embedding. Cross-sections (6 μm-thick) were obtained and submitted to the labelling of pRB1, Ki67, PCNA, NANOG and Casp3. The sections were dewaxed in xylene, hydrated and submitted to heat antigen retrieval using citrate buffer (pH 6.0) for 10 minutes. The slides were treated with 5% BSA and incubated with the primary antibodies pRB1 (1:200, Cell Signal, USA), Ki67 (1:200, Abcam, USA), PCNA (1:200, Biocare Medical), NANOG (1:200, Abcam, USA) and Casp3 (1:250, Cell Signal, USA) overnight at 4° C. The slides were washed in PBS (0.05 M, pH 7.2) 3x and incubated with the secondary antibody (DAKO Detection System - K0690, USA). The slides were washed again in PBS and then incubated with the streptavidin-peroxidase (DAKO Detection System - K0690, USA). The reactions were revealed with DAB (K3468, DAKO, USA) and the nuclei were stained with Harris Hematoxilyn. Negative controls (primary antibody omission) were performed for each antibody. The slides were carefully analysed and the positive germ cells were counted using the Leica Analysis System (LAS) (Cambridge, England). For each embryonic and postnatal age, 6 individuals were analysed (n=6). The mean number of positive gonocytes was converted to percentage for better representation and interpretation of the results.

### Gonocyte Isolation and Casp3 Inhibition

The testes from ten to twelve 19dpc embryos form 9 different mothers were collected and dissociated using collagenase/dispase (Roche, 1mg/ml), at 37°C, for 45 min associated with mechanical dissociation using a P1000 pipette. The suspension was placed in 6-well culture plates, in a proportion of two testis per well, in DMEM supplemented with 10% FBS and 1% penicillin/streptomycin, at 37°C and 5% de CO_2_. The whole testis suspension was maintained for 4h at this condition. Since the somatic cells from the gonad, especially Sertoli cells and the fibroblasts, adhere very easily to the bottom of the plates and the gonocytes do not adhere at all, this protocol was used to separate the gonocytes from the somatic cells. The plates form the first incubation (4h) were analysed in AZ100 macroscope (Nikon) to check whether the somatic cells were attached. To check whether the supernatant contained significant contamination with somatic cells and whether gonocytes could have been dragged to the bottom of the plate, the supernatant was investigated for the expression of Sertoli cell and interstitial cell markers using primers for *Sox9* and *Gata4*, and the attached cells were investigated for the expression of the germ cell markers *Dazl* and *Mvh* (not shown). None of these markers were detected, indicating non-relevant contamination of somatic cells in the supernatant or of germ cell retention among the attached cells.

Subsequently, then, the supernatant containing the germ cells (gonocytes) was collected and transferred to new 6-well plates in the same culture medium with the addition of 10 µM of the Casp3 inhibitor (Ac-DMQD-CHO, Cat. 235421 – Calbiochem). The culture was maintained for more 20h. The dose of the Casp3 inhibitor was chosen based in the literature [24] and on previous test experiments form our group. These experiments were performed twice in biological triplicate with control experiments (absence of Casp3 inhibitor) running in parallel in each experiment.

### Flow Cytometry

Gonocytes viability was determined by staining with propidium iodide (10µg/mL - BD Bioscience). The cells were incubated in the dark, at room temperature, for 10 min, following the manufacturer’s protocols. After incubation, cells were analysed in a FACS CANTO II (BD Biosciences) flow cytometer. A total of 20.000 events were analysed and plotted in a SSC-HxSSC-A dot plot to exclude doublets. From a single cell gate, debris was depleted by a region gate determined in a FSC-AxSSC-A dot plot. After this, the percentage of propidium iodide positive cells was determined based on the fluorescence of non-staining cells. This experiment was performed in the Laboratory of Lymphocyte Ontogeny at Department of Microbiology, Immunology and Parasitology at UNIFESP/São Paulo.

## RT-PCR

The total RNA was isolated from embryo gonads at 15, 17 and 19dpc and from the gonocyte suspension using TRIzol regent (Invitrogen). The cDNA was obtained using the Superscript IV Reverse Transcriptase (Invitrogen). Quantitative RT-PCR was performed using PowerUp Sybr Green Master Mix (Thermo Fisher Scientific) and specific primers for *Casp3, Pcna, Nanog, Sox2, p21cip, p27kip, Bcl2* and *Bax* for the gonocytes and *Sox2* and *Nanog* for the 17dpc and 19dpc gonads, in a Roche LightCycler96 platform. The primer sequences are described in Table 1.

**Table 1:**
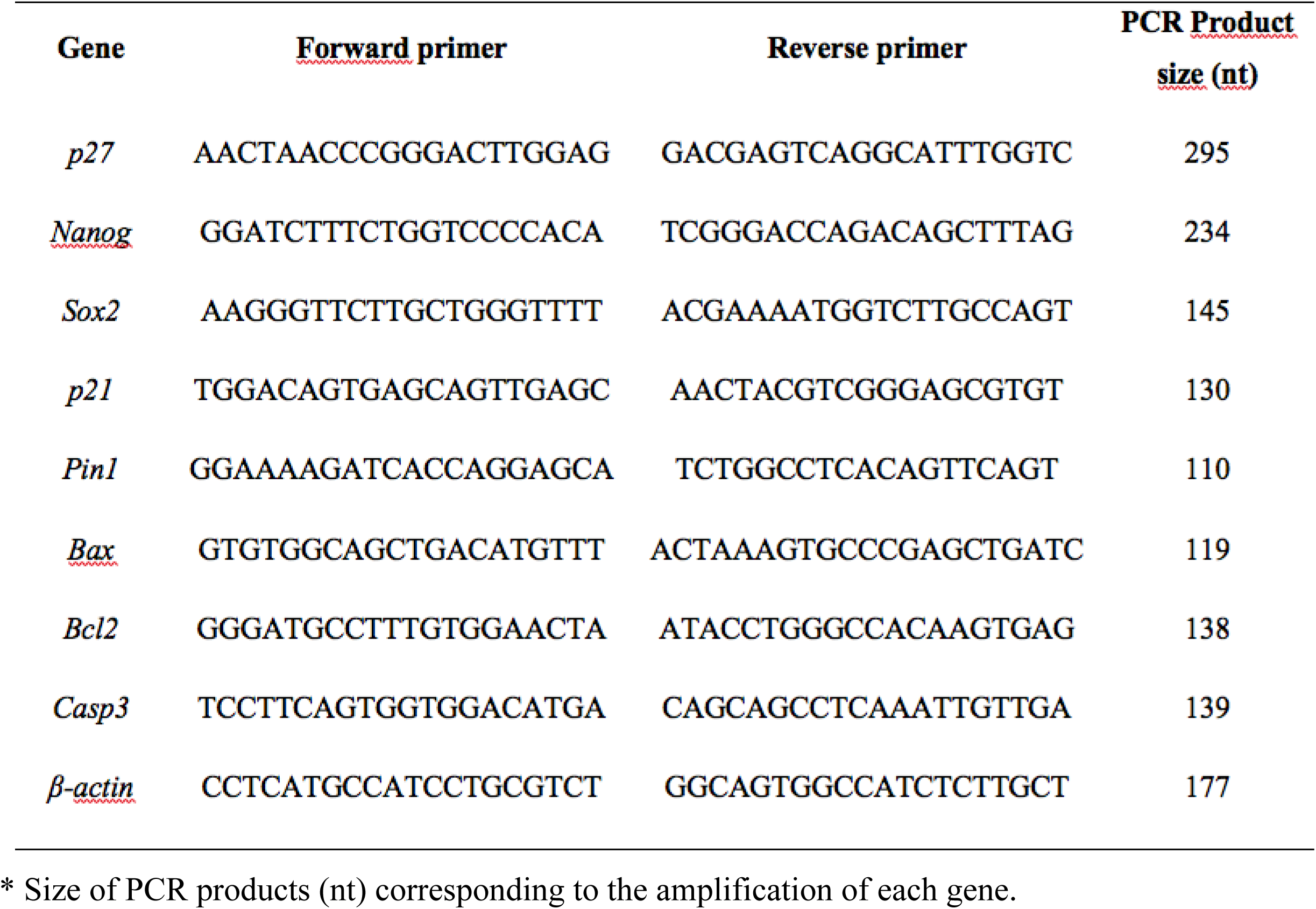
Sequences of primers used for RT-qPCR and the size of PCR products corresponding to the amplification of each gene.

### Western Blot

The total protein extract from ten testes at 19dpc was heated as instructed by the manufacturer (Bolt™ LDS Sample Buffer and Bolt™ Reducing Agent - Thermo Fisher Scientific). After SDS-PAGE (Bolt™ 4-12% Bis-Tris Plus Gel - Thermo Fisher Scientific), proteins were transferred to nitrocellulose membrane (Bio-rad). After the transference, the membrane was incubated with blocking solution (Western Dot 625^®^ - Cat. W10142, Invitrogen) for 1h and then with anti-NANOG primary antibody (1:1000, Abcam, USA) overnight at 4°C and shaking. The membrane was washed in Washing Buffer (Western Dot 625^®^ - Cat. W10142, Invitrogen) and incubated with the biotinylated secondary antibody (anti-rabbit Western Dot 625^®^ - Cat. W10142, Invitrogen) for 1h. The membrane was then washed and incubated with QDot625 conjugated with streptavidin (Western Dot 625^®^ - Cat. W10142, Invitrogen).

### Statistical Analysis

The number of gonocytes obtained from the IHC and from viability experiments were analysed using one-way ANOVA test. Data from RT-qPCR was analysed using Student’s *t* test. The differences were considered significant when p≤0.05.

### Ethical Approval

This study was carried out according to the rules of the local committee for animal care (approval number: 9806200114).

## Results

### Ki67, pRB1, p27^kip^ labelling

The labelling of the proliferation marker Ki67 (Figs 1A-1C) and of cell cycle regulator pRB1 (Figs 1D-1F) indicates that the rat male germ cells start to enter quiescence around 17dpc and that at 19dpc all gonocytes are quiescent. The percentages of Ki67- and pRB1-positives cells at each age decreased as age increased (Fig 2). At 15dpc (Fig 1A) 98% of the gonocytes were ki67-positive, indicating that they are proliferating. At 17dpc (Fig 1B), 40% of the gonocytes were Ki67-positive, what suggest that the quiescence process has initiated. At 19dpc (Fig 1C) no gonocytes were positive for Ki67, confirming that they are quiescent at this stage. Sertoli cells were negative at 15dpc (Fig 1A). At 17dpc there were positive and negative Sertoli cells (Fig 1B), whereas at 19dpc (Fig 1C) all Sertoli cells were Ki67-positive.

**Fig 1.**
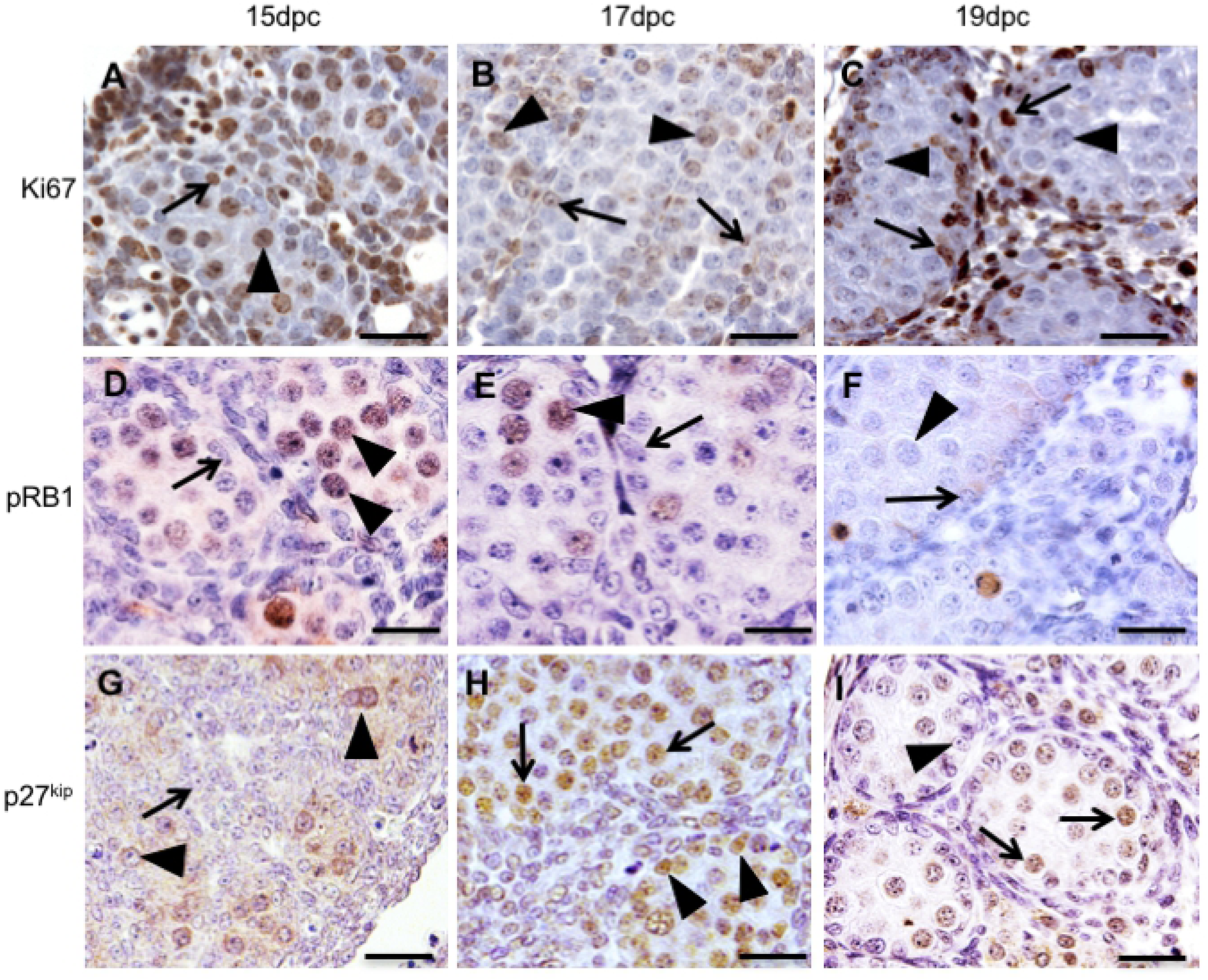
Immuno-labelling of Ki67 (Figs 1A, B, C), pRB1 (Figs 1D, E, F) and p27^kip^ (Figs 1G, H, I) in the gonads of 15, 17 and 19dpc rat embryos. Ki67 and pRB1 were present in the gonocytes (arrowheads) at 15 and 17dpc, but not at 19dpc, indicating that these cells are not cycling at this age. Sertoli cells (arrows) were positive for Ki67 and negative for pRB1 at the three ages. P27^kip^ was detected in the cytoplasm of gonocytes (arrowheads) at 15dpc and in their nucleus at 17 and 19dpc. Sertoli cells (arrows) were negative for p27kip at 15dpc and 19dpc, but showed nuclear labelling at 17dpc (Scale bar = 33 µm)

**Fig 2.**
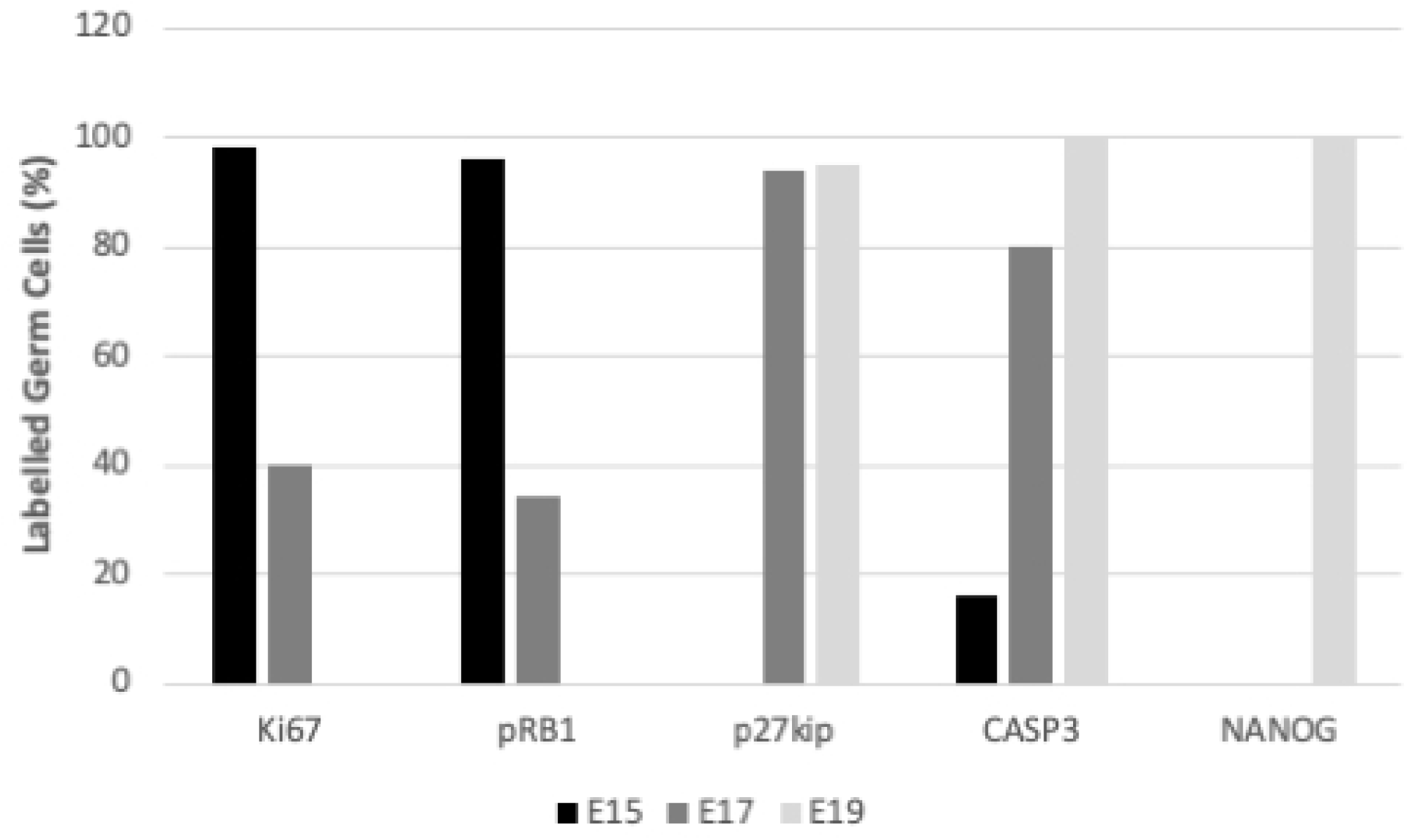
Percentage of Ki67-, pRB1-, ASP3- and NANOG-positive gonocytes in the gonads of 15, 17 and 19dpc rat embryos. The number of Casp3-positive gonocytes increased from 15dpc to 19dpc whereas the number of Ki67- and pRB1-positive gonocytes decreased.

pRB1-positive gonocytes were observed at 15dpc (Fig. 1D) and 17dpc (Fig. 1E). The labelling was observed in nuclei of the gonocytes. pRB1 was not detected in gonocytes at 19dpc (Fig 1F). The number of pRB1-positive gonocytes (Fig 2) was higher at 15dpc (96%) than at 17dpc (34%) and was reduced to zero (0%) at 19dpc. Sertoli cells were negative for pRB1 at the three ages.

The labelling of p27^kip^ was present in the cytoplasm of germ cells at 15dpc with no staining in the nucleus (Fig 1G), whereas at 17dpc (Fig 1H) and 19dpc (Fig 1I) this protein was detected in the nucleus of 94% and 95% of germ cells, respectively (Fig 2). Sertoli cells were negative for p27kip at 15dpc (Fig 1H) and 19dpc (Fig 1I), but showed nuclear labelling at 17dpc (Fig 1G).

### CASP3 and NANOG detection

CASP3 detection (Figs 3A-3C) showed an inverse pattern of that observed for Ki67 and pRB1 from 15dpc to 19dpc. At 15dpc (Fig 3A) only 16% of the gonocytes and some apoptotic bodies CASP3-positive. At 17dpc (Fig 3B) the number of CASP3-postive gonocytes increased and 80% were positive for this protein. At 19dpc (Fig 3C) 100% of these cells were positive for CASP3. Sertoli cells were negative at all ages. A similar labelling pattern was observed for NANOG. Very rare labelling (0%) was observed at 15dpc (Fig 3D) and 17dpc (Figs 2 and 3E). On the other hand, at 19dpc intense and abundant NANOG labelling was observed in the cytoplasm of all gonocytes (Figs 2 and 3F). NANOG labelling at 19dpc was very similar to CASP3 labelling (Figs 3C and 3F).

**Fig 3.**
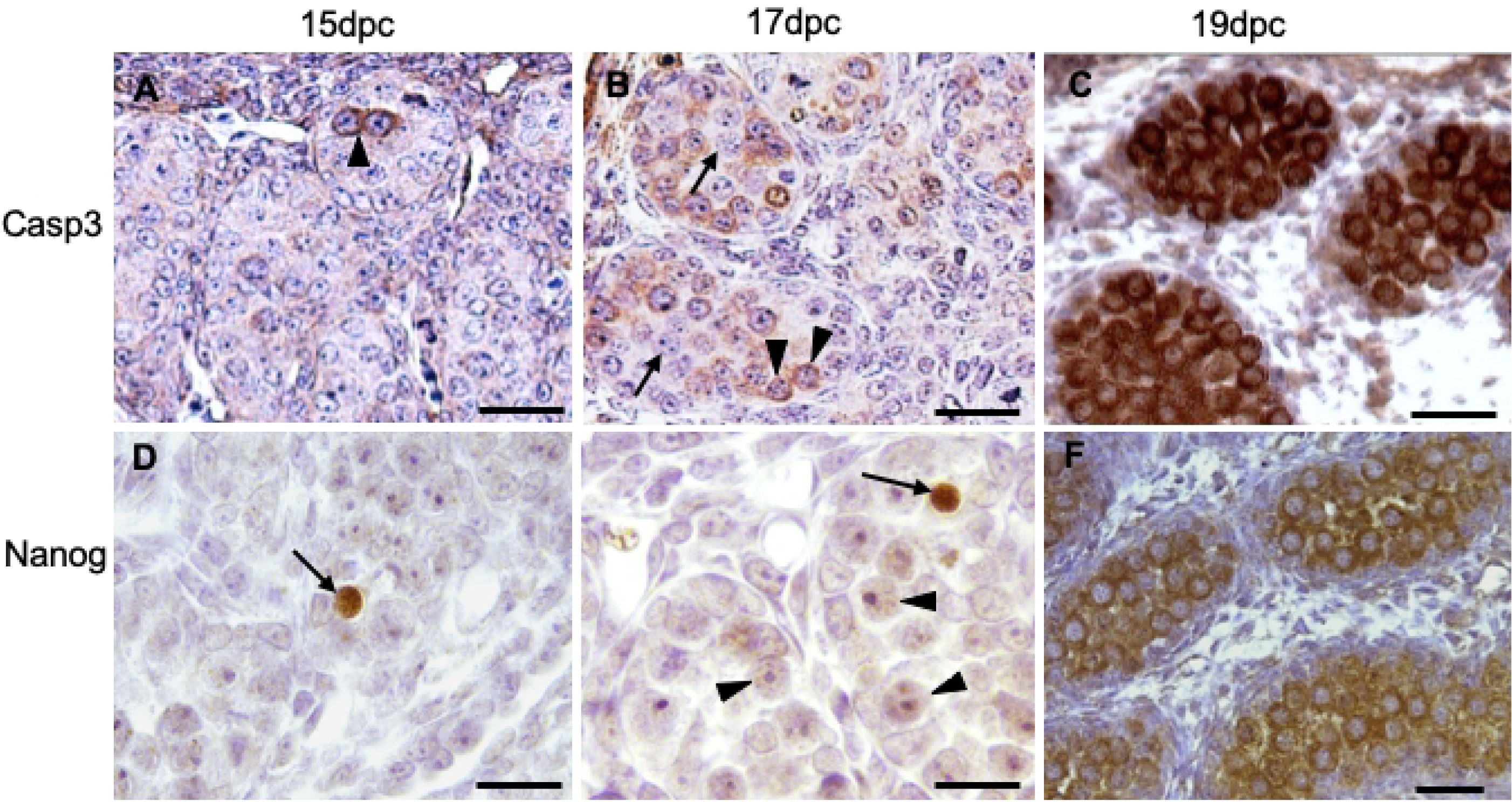
Immuno-labelling of CASP3 and NANOG in in the gonads of 15, 17 and 19dpc rat embryos. CASP3 labelling (arrowheads) is rare at 15dpc (Fig 3A) and increased at 17dpc (Fig 3B). CASP3-negative germ cells are also observed (arrows). At 19dpc (Fig 3C) all gonocytes are positive for CASP3. NANOG labelling (arrows) is very rare at 15dpc (Fig 3D) and 17dpc (Fig 3E), although at 17dpc a weak labelling is seen in most germ cells (arrowheads). As observed for CASP3, at 19dpc (Fig 3F) all gonocytes are positive for NANOG. (Scale bar = 33 µm)

The presence of cleaved CASP3 was confirmed in the 19dpc testis by Western blot (Fig 4) but not at 15dpc and 17dpc probably due to the low expression levels at these ages. However, despite the intense and abundant labelling of NANOG in the immunohistochemistry assay and the detection of *Nanog* expression by RT-qPCR, the presence of this protein could not be confirmed by Western blot analysis.

**Fig 4.**
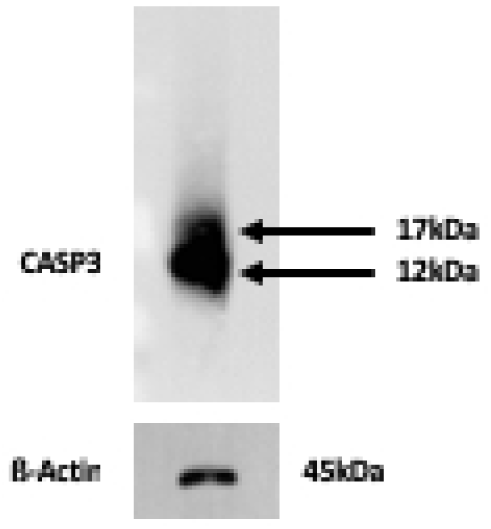
Caspase-3 expression on 19dpc testis. Two bands of 17kDa and 12kDa (arrows) are observed and correspond to the two subunits of cleaved CASP3. ß-ACTIN was used as reference.

### Gonocyte purification and Viability

To investigate the possible functions of CASP3 during gonocyte quiescence, these cells were isolated and incubated with CASP3 inhibitor. Because some of the germ cell markers present in the membrane, such as SSEA1 and DBA (*Dolichus biflorus* agglutinin), are not present at this phase of development, we decided to use a different method of purification. Based in previous observations that quiescent gonocytes do not adhere to treated culture plates or even to feeder cells such as fibroblasts or Sertoli cells (personal observations), we incubated 19dpc dissociated gonads in pre-treated culture plates for 4h. We observed that the somatic cells adhere to the bottom of the plate (S1 Fig), whereas the gonocytes stay in the suspension. So, the suspension was collected and transferred to new treated plates where the incubation with the CASP3 inhibitor for 20h took place. The RT-PCR analysis showed that germ cell markers *Dazl* and *Mvh* were not detected in the attached cells but were detected in the suspension. Conversely, the Sertoli cell markers *Gata4* and *Sox9* were not detected in the suspension but were detected in the attached cells.

To check whether the cells were viable after the end of the incubation period, their viability was tested by flow cytometry. We observed that in the control cultures gonocyte viability was higher (53%) than in the cultures treated with CASP3 inhibitor (26,1%) (Fig 5).

**Fig 5.**
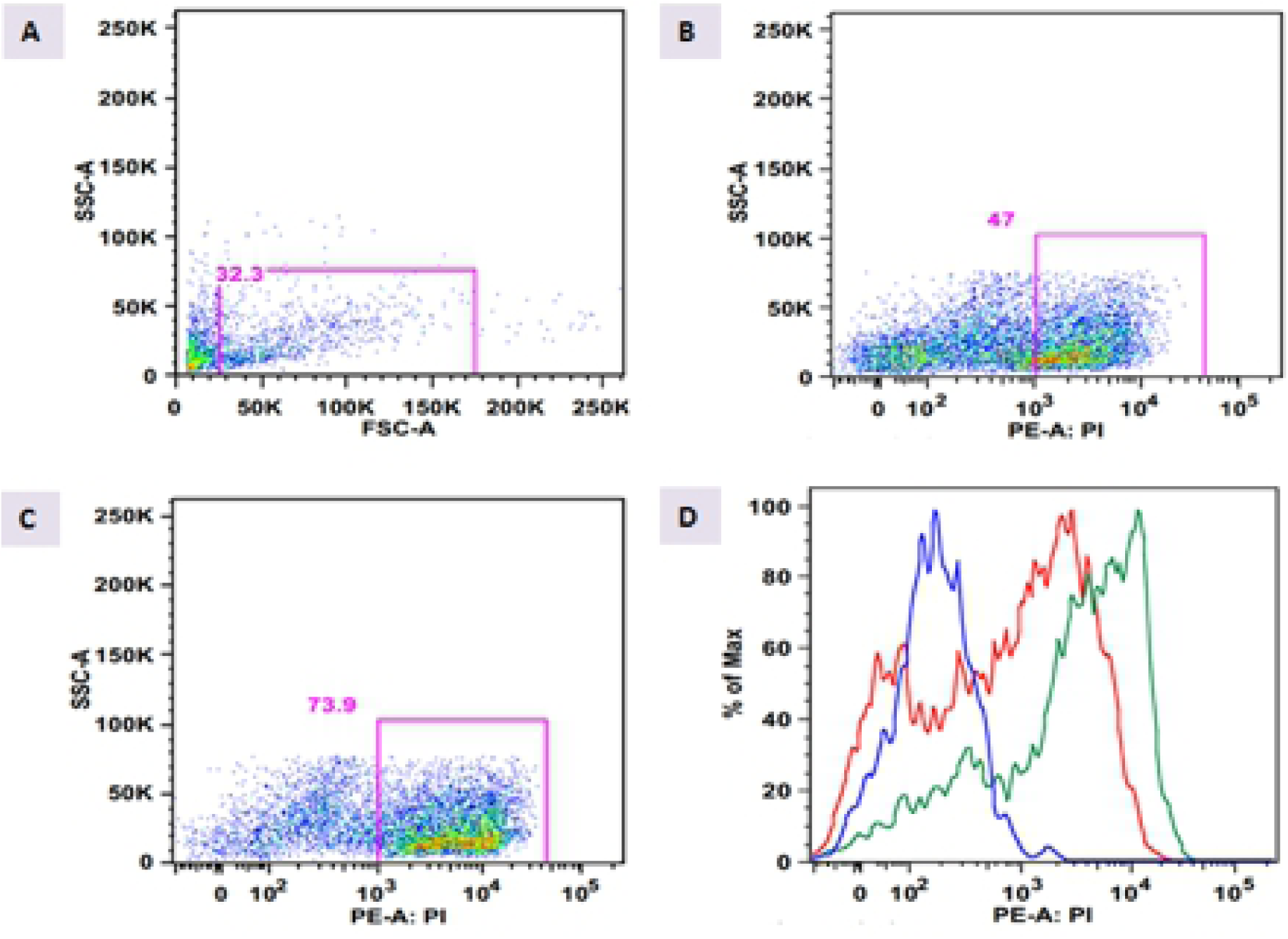
Representation of Flow cytometry of gonocytes submitted to culture. A: non-labelled sample. The pink rectangle indicates the population of interest (gonocytes). B and C: Control and treated cultures, respectively. The pink square indicates the gonocytes labelled with propidium iodide (PI). D: Diagram showing the peaks corresponding to PI labelling. Blue: non-labelled sample; Red: control culture; Green: treated culture.

### Expression of Cell cycle, Apoptosis and Pluripotency Genes

The expression of the cell cycle, apoptosis and pluripotency genes *Pcna, p27*^*kip*^, *Kras, p21*^*cip*^, *Casp3* and *Pin1* was investigated after CASP3 inhibition *in vitro*. In consonance with our hypothesis, CASP3 inhibition lead to a decrease of *p27*^*kip*^ and *Bcl2* expression and to an increase of *Pcna* expression when the ∆Cq was considered (Table 2). When the calibrator sample (E19) was included and the fold change was used as a parameter, the expression of *p21*^*cip*^, *Bcl2* and *Casp3* showed an important decrease (50% or more) in the treated gonocytes when compared with the control gonocytes, whereas the expression of *Pcna* showed an increase (70%) in the treated gonocytes (Table 2).

**Table 2:** Expression of cell cycle and apoptosis markers presented as ΔCq and fold change values.

| Gene | $\Delta Cq$ CC | $\Delta Cq$ TC | p value | Fold change CC | Fold change TC | p value |
| --- | --- | --- | --- | --- | --- | --- |
| *Pin1* | $6.82 \pm 0.65$ | $7.083 \pm 1.47$ | 0.73 | 0.0442 | 0.0405 | 0.847 |
| *p21* | $4.2 \pm 2.7$ | $3.67 \pm 0.09$ | 0.322 | 28.928 | 11.976 | 0.294 |
| *Bcl2* | $6.02 \pm 0.48$ | $8.83 \pm 1.23$ | 0.0478* | 0.303 | 0.0499 | 0.039 |
| *Bax* | $2.343 \pm 0.58$ | $2.73 \pm 0.14$ | 0.329 | 1.253 | 0.911 | 0.31 |
| *Casp3* | $4.977 \pm 0.34$ | $6.03 \pm 0.83$ | 0.114 | 0.153 | 0.0798 | 0.039 |
| *p27* | $4.645 \pm 0.64$ | $7.73 \pm 0.10$ | 0.0216* | 6.478 | 0.728* | 0.040 |
| *Pcna* | $5,423 \pm 0.78$ | $4,65 \pm 0.25$ | 0,0267* | 0,103 | 0,177* | 0.03 |

## Discussion

The control of germ cell proliferation and differentiation during the embryonic phase is fundamental for the normal development of these cells. Although the existence of a period of mitotic arrest during embryonic and early postnatal germ cell development in rats has been mentioned [6,25], the precise age when this occurs is not known. In the present study we show that the first signs of quiescence in rat gonocytes are observed at 15dpc and that at 19dpc all gonocytes are quiescent. We also strongly suggest that CASP3 plays a role in this process and describe some of the genes and proteins possibly involved.

It has been shown that CASP3 has non-apoptotic functions such as in the cell cycle, controlling proliferation [22, 26] and in cell differentiation [27]. The administration of CASP3 inhibitors affects the proliferation of T cells [28, 29] and the differentiation of skeletal muscle differentiation [27].

In the present study, the presence of the cell cycle markers Ki67, pRB1 and p27^kip^ was investigated during rat germ cell proliferation and quiescence and compared with CASP3 detection. Ki67 and pRb1 labelling reduced in germ cell nuclei as they entered quiescence, showing an inverse pattern of that observed for CASP3, i.e., the number of CASP3-positive gonocytes increased with the embryo age while the number of pRb1- and Ki67-positive gonocytes decreased. While Ki67 is an ordinary proliferation marker, the role of pRb1 in gonocyte quiescence has been demonstrated in mice [14, 23]. Rb1 is a potent regulator of the cell cycle, acting in proliferation, differentiation and apoptosis. In its hyperphosphorilated form, Rb1 binds to transcription factor E2F, inhibiting the cell cycle progression. Western et al. (2008) showed that the Rb1 expression is downregulated in gonocytes as they enter quiescence. This data agrees with those obtained in the present study. Similarly to CASP3 immunolabelling, the presence of p27^kip^, a cyclin inhibitor, in germ cell nuclei was associated with quiescence entry. Here p27^kip^ was detected in the cytoplasm of proliferating germ cells and, as they entered quiescence, its detection became nuclear. The increase of p27^kip^ expression in mouse gonocytes as they enter quiescence has been reported [14, 23]. Interestingly, p27^kip^ is a substrate of CASP3 [30]; thus, the nuclear localization of p27^kip^ might be necessary not only for its function but also to avoid degradation by CASP3.

The labelling pattern of CASP3 compared to Ki67, pRb1 and p27^kip^ labelling raises the hypothesis that CASP3 has a role in rat gonocyte quiescence. Thus, we decided to inhibit CASP3 in gonocytes and look at the expression of cell cycle markers that are substrates of CASP3 (*p27*^*kip*^ and *p21*^*cip*^) at mRNA level. Although the investigation of the same markers at protein level could provide relevant data to this study, this was not feasible due to the low amounts of cells available. Importantly, the data obtained was in consonance with our hypothesis, since we observed a decrease in *p27*^*kip*^ and *p21*^*cip*^ expression in gonocytes after CASP3 inhibition.

As previously mentioned, the existence of the quiescence period seems to be essential for the normal differentiation of male germ cells in rats and mice, although the exact role of this phenomenon has not been elucidated yet. In other cell types, it is not known whether the quiescence is a passive process in which the cells keep stationary or whether it is an active process necessary to keep cell integrity. The quiescent period begins from the G1 checkpoint, when the cell must decide whether to continue the cell cycle or enter a rest phase, when DNA integrity and the cell cycle machinery is checked [31]. This raises the hypothesis that the abundance of CASP3 in quiescent gonocytes could somehow be related to the cell cycle checkpoint, although further experiments are necessary to confirm this hypothesis. PCNA is one of the proteins that act during this checkpoint. Its interaction with other proteins responsible for the cell cycle control, such as p21, cyclins and cyclin-depent kinases (CDK), has been described [32]. Although PCNA is widely used as a proliferation marker, in germ cells it plays an important role in DNA repair [33]. Here, the inhibition of CASP3 lead to an increase of *Pcna* expression in gonocytes. Thus, we decided to look at the expression of PCNA in quiescent gonocytes and showed that it is indeed detected in these cells (S2 Fig), confirming that PCNA might not be an ideal marker for germ cell proliferation and might function as a cell cycle regulator in quiescent rat gonocytes. So, the increase of *Pcna* expression observed after CASP3 inhibition could be rather related to the prevention of DNA damage than to an attempt to resume proliferation.

The negative relationship of the pluripotency marker OCT4 with male germ cell arrest has been shown in mice [14]. In rats we have previously shown that OCT4 is detected in germ cells from 13dpc to 17dpc [34], is absent at 19dpc and comes back at 8dpn [10], i.e., also in rat germ cells this pluripotency marker is absent in quiescent gonocytes. It has been suggested that the downregulation of *Oct4* is necessary for normal germ cell differentiation and that an inefficiency of *Oct4* downregulation can be associated with the development of testicular germ cell tumour (TGCT) in humans [35-37]. In rats, the occurrence of TGCT has not been described so far, even after the use of substances that are potential tumour inducers, such as the phthalates [38]. It is important to mention that gonocytes do not enter an effective period of quiescence in humans. Thus, one of the hypotheses to explain this is that the quiescence period can be a key event in the prevention of TGCT. In other cell types, the quiescence period works as tumour suppressor [39, 40]. If the gonocyte quiescence is considered a standard process of checkpoint decision, it can be suggested that during this period the male germ cells ensure that possible DNA damages are fixed. Although we and others have observed a negative relation between pluripotency markers and gonocyte quiescence, here we show that the expression of Nanog, another important pluripotency marker, was detected in quiescent gonocytes at both RNA and protein levels. Although we were not able to confirm NANOG expression by Western blot assay, the pattern of NANOG immuno-labelling in quiescent gonocytes was identical to that observed for CASP3. NANOG is a typical marker for embryonic stem cells (ES) [41, 42] and its downregulation is associated with ES differentiation [43]. Fujita et al. (2008) showed that CASP3 cleaves NANOG and mediates ES differentiation [44]. According to Zhang et al. (2009) NANOG controls G1 to S transition in ES by regulating CDK6 and CDC25A at the transcriptional level [45]. Taken together the similarity between CASP3 and NANOG localization in quiescent gonocytes and the fact that CASP3 cleaves NANOG, we suggest that these two proteins might work together during rat gonocyte entry into mitotic arrest.

In conclusion, this study shows that the first signs of male germ cells quiescence in the rat appear at 15dpc and coincides with the onset of CASP3 detection, leading us to suggest that CASP3 might have a role in this phenomenon. Furthermore, we suggest that Nanog and Pcna could be CASP3 partners. Further studies are necessary to confirm whether CASP3 regulates male germ cell cycle and, if so, how it occurs.

## Funding

The authors thank Coordenação de Aperfeiçoamento de Pessoal de Nível Superior - Brasil (CAPES) - Finance Code 001 and Fundação de Amparo à Pesquisa do Estado de São Paulo (FAPESP) for research funding (Proc. Nr. 2012/08951-1) and for student grants (Proc. Nr. 2012/02400-3).

## Conflict of Interest

The authors declare no conflict of interest

## Supporting information

**S1 Fig.**
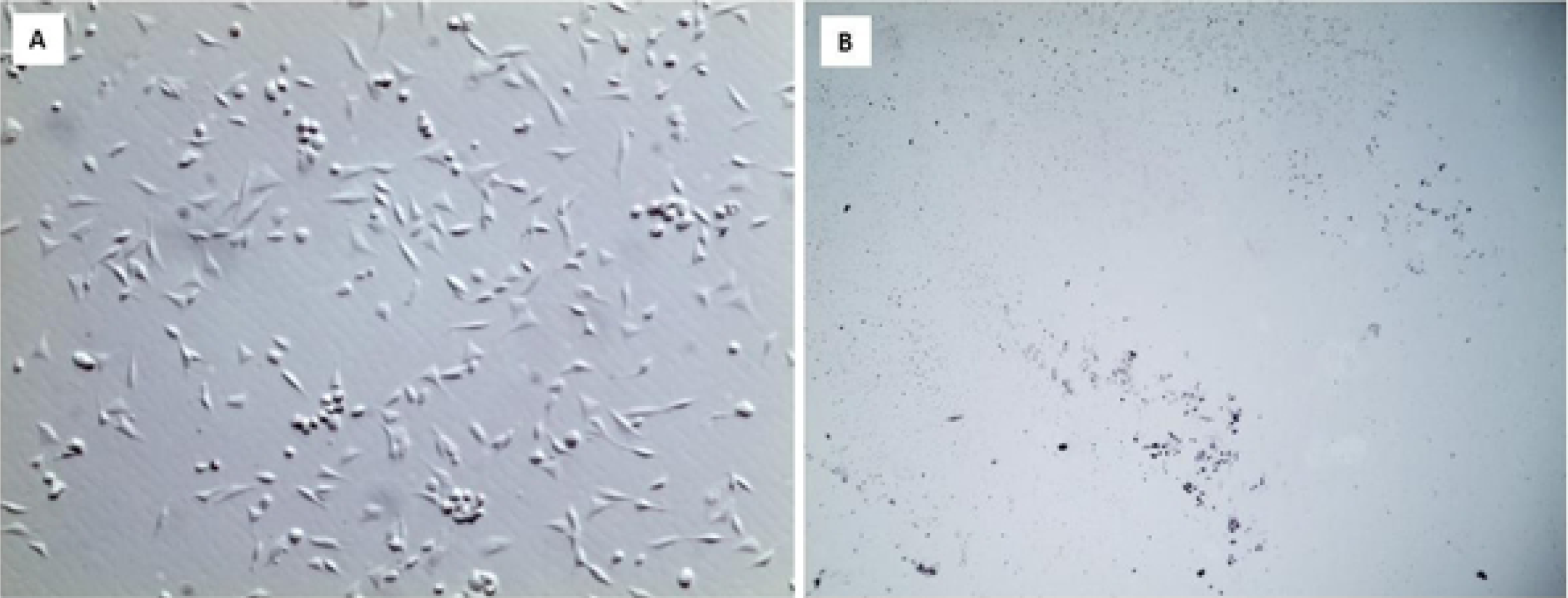
Photomicrographs of the bottom of the culture plates after 4h (A) of incubation of 19dpc dissociated (whole) testis and after 20h of incubation of the gonocyte suspension. In A the somatic cells, especially Sertoli cells and fibroblasts, are attached. This photograph represents the bottom of the plate after the gonocyte suspension was removed. In B no cells are observed in the bottom of the plate after the gonocyte suspension was incubated for 20h and then removed.

**S2 Fig.**
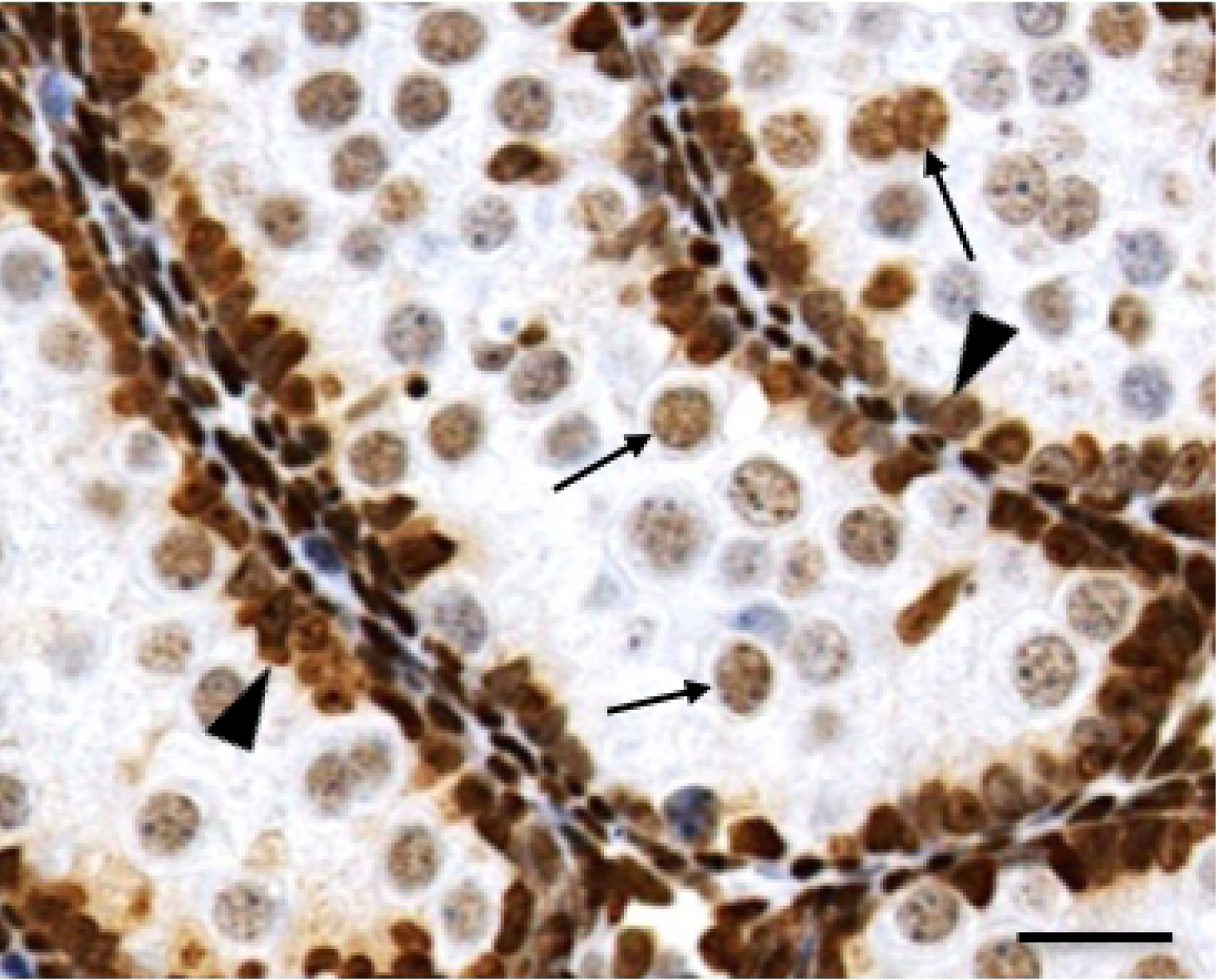
PCNA immune-labelling in 19dpc testis. PCNA was detected in the nucleus of gonocytes (arrows) and Sertoli cells (arrowheads). Scale bar: 32µm.

